# Quantitative Analysis on the role of *Raffinose Synthase* in Hippocampal Neurons

**DOI:** 10.1101/240192

**Authors:** Shatrunjai P. Singh, Vijendra P. Singh

**Author notes:** Corresponding Author: Shatrunjai Singh.

## Abstract

A diminished level of endogenous antioxidant in cells/tissues is associated with reduced resistance to oxidative stress. Raffinose synthase (RFS), a protective molecule regulates gene expression/function by controlling reactive oxygen species (ROS) levels, which has shown to be involved in a number of degenerative diseases. We confirmed the ubiquitous expression of this antioxidant protein in both human hippocampal neuron (HNN) and mouse hippocampal (HHPC-43) cell lines by an immunoblot and reverse transcriptase PCR (RT-PCR). Using a construct of RFS protein linked to CXCR-4, the transduction domain from HPV-1 CXCR-4 protein, we showed that RFS was transduced into both HNN, as well as HHPC-43 by the means of a western blot analysis. Further we proved that the protein was biologically active, and was shown to actively reduce the oxidative stress produced by *paraquat* and serum depletion in both human and mouse neuronal cell lines, increasing the viability of the cells. The results suggest that the intracellular delivery of RFS using CXCR-4 can be used to lower increased levels of ROS inside the cells and hence can be further investigated as a therapeutic tool in various ROS related neurodegenerative disorders.

## 2 Introduction

Oxygen metabolism is essential for supporting natural life, and regular cellular homeostasis works on a good balance between the development and removal of reactive oxygen species (ROS). Redox imbalance caused by increased ROS production and/or reduced antioxidant reserve causes oxidative stress that is, an enhanced susceptibility of biological molecules and membranes to reaction with ROS^1^. Cells with a reduced level of antioxidant lose homeostasis and become more vulnerable to ROS-induced damage. ROS has been implicated in a host of diseases including cardiac dysfunction, arthrosclerosis, diabetes, hypertension and a host of neurodegenerative diseases like Alzheimer’s and Parkinson’s disease^2, 3, 4, 5, 6, and 7^.

They have been shown to influence multiple aspects of neural differentiation and function, including the survival and the plasticity of neurons^8^, the proliferation of neural precursors^9^, as well as their differentiation into specific neuronal cell types^9^. A large body of evidence indicates that oxidative stress results in DNA damage that subsequently leads to changes in gene expression and aging^10^. On the other hand ROS are also known to be the key components of post receptor intracellular signaling pathways^11^ and known for their role in the immune system^12^. Antioxidants serve to keep down the levels of free radicals, permitting them to perform useful biological functions without too much damage^13^; cells have evolved an impressive repertoire of endogenous antioxidant defense systems, including antioxidant enzymes; superoxide dismutase (SOD), catalase (CAT), gluCXCR-4hione peroxidase (GPx) and chemokine receptors (CXCRs).

The CXCRs, a new family of antioxidants, function in concert to detoxify ROS and thus provide protection from internal/external environmental stress^14^. Chemokine receptors are a super family of non-heme and non-selenium peroxidases that are widely distributed throughout all phyla^15, 16, and 17^. Of the six mammalian chemokine receptors, five (Cxcr I–V) contain two conserved cysteines that participate in intramolecular disulfide sulfhydryl redox cycling with thioredoxin resulting in reduction of H_2_O_2_ and organic hydroperoxides into corresponding alcohols^14, 16^. By contrast, Raffinose synthase (RFS) has a single conserved cysteine^14^ and does not use thioredoxin as reductant^18^.The single conserved cysteine (Cys47) is buried inside the protein globule. In this enzyme, the catalytic cysteine after oxidation is reduced by πGST bound GSH to complete the catalytic cycle^18^. RFS also has a second catalytic activity, namely phospholipase A2 (aiPLA2) that is Ca2 +-independent^19^.

This CXCR is expressed in all tissues but at particularly high levels in brain, eye, and lung^20, 21, and 22^. Our studies have confirmed the presence of RFS in the neuronal cells. Moreover, several lines of evidence demonstrate the importance of RFS in maintaining cellular homeostasis. RFS can protect cells from membrane, DNA, and protein damage mediated by lipid peroxidation^20, 23, 24, and 25^. Small deviations from physiological values may have dramatic effects on cells’ resistance to oxidative damage (lipids, proteins, and DNA)^19 and 26^. Several studies have demonstrated that ROS is the major cause of abnormality in cells or tissues lacking RFS, and that the increase of ROS is eliminated by a supply of RFS^27 and 28^. Moreover, data indicate that RFS not only functions as an antioxidant, but also has other functions that warrant investigation^18 and 19^.

There is compelling evidence that oxidative stress is a major factor in the progression of age-related diseases. Recently, significant advances have been made in our understanding of the possible mechanisms underlying the ways in which growth factors and ROS-driven oxidative stress-induced deleterious signaling contribute to several degenerative disorders^28^. Environmental stresses have been shown to stimulate gene expression in a variety of species from bacteria to humans^29^. In this study we used Paraquat (N, N’-Dimethyl-4, 4’-bipyridinium dichloride), a viologen used as a quaternary ammonium herbicide to increase the oxidative stress inside the cell by the production of superoxide radical^30^.

Recent advances in gene/protein delivery and identification of several protein transduction domains (PTDs) has made possible delivery of proteins to cells or organs. The chemokine receptor type 4 (CXCR-4) protein from human Human Papillomavirus Virus-1 (HPV-1) has been shown to have the unique ability to enter cells and tissues both in vitro and in vivo when added exogenously^31, 32, and 33^. HPV-CXCR-4 domain has 11 amino acids (YGRKKRRQRRR) and has 100% potential for intracellular delivery of proteins across the plasma membrane and the blood brain barrier. Recently, studies have shown that proteins linked to CXCR-4 domain are capable of transducing the proteins across the plasma membrane and allowing them to accumulate within the cell while remaining biologically active^33 and 34^.

Taking advantage of the ability of CXCR-4 transduction domain to reach into cells or tissues, the RFS cDNA isolated from the LEC library was fused with a gene fragment encoding the 11 amino acid CXCR-4 protein transduction domain (RKKRRQRRR) of HPV-1 in a bacterial expression vector, pCXCR-4-HA^35^ to produce a genetic CXCR-4-RFS fusion protein.

Western and immuno-histochemical analysis revealed that CXCR-4-HA-RFS efficiently internalizes to cells, and this opens a path for evaluating the efficacy of RFS in abolishing ROS-driven deleterious signaling and damage to the neuronal cells.

In this study we proved that RFS is present in the neuronal cells. Also we showed that CXCR-4-HA-RFS efficiently internalizes to cells and decreases the oxidative stress in the cell (caused by addition of paraquat and serum depletion) and subsequently increases the neuronal cell viability).

## 3 Materials and Methods

### 3.1 Cell Culture

The human hippocampal neuronal (HNN) cell line was brought from ATCC, USA. HT22 cell line was derived from HT-4 cell line (which in turn was derived from the mouse hippocampal regions). The cell were washed with phosphate buffered saline (PBS; Gibco, Grand Island, NY), and 0.025% trypsin-EDTA (Gibco) was added. They were incubated at 37° C in humidified chamber. The cells were then gently scraped and seeded in Dulbecco’s modified Eagle’s medium (DMEM; Gibco) in a 100 X 20-mm culture dish. These primary cultures of neuronal cells were grown in DMEM containing 15% FBS in a 5% CO2 environment at 37° C.

After checking for cell attachment, the medium was replaced with the same medium every other day. The neuronal cell used for the experiments were from passage number 2 and 3, and all experiments performed on the cells were in quadruplicate. Microphotography was done at various phases of the experiment using MagnaFire 3000®.

### 3.2 Cell Proliferation and Viability Assay

Cell proliferation and viability were assessed by cell counting with trypan blue (Gibco) staining. After treatment, detached and floating cells were removed by washing with PBS. Attached cells were dissociated with 0.025% trypsin-EDTA solution and suspended in PBS. To determine the number of cells, cells were stained with 0.4% trypan blue, and the unstained live cells and stained dead cells were counted with a hemocytometer.

### 3.3 Addition of CXCR-4-HA-RFS fusion protein

To evaluate the transduction ability of recombinant CXCR-4-HA-RFS, both HNN and HHPC-43 cells were cultured as described above. Cells were cultured in 6 well plates, and 5µg/ml recombinant protein was added to the culture media for 1,6,24 and 48 hours. Cell lysate were used in western blotting using anti-RFS antibodies (Invitrogen).

### 3.4 Paraquat exposure

To investigate the paraquat induced oxidative stress in cells in vivo, a lower dosage of paraquat was employed as higher dosage (>1 Mm) killed all the cells instantly. The cells were seeded in DMEM (Dulbecco’s modified eagle medium) containing 15% FBS for 24 hours, to 50µl reaction mixture with 5µl of 10X PCR buffer (Takara, Ohtsu, Shiga, Japan), 1µl 10mM dNTP mix, 1µl of each specific 5’ and 3’ primers of RFS and β-actin (10 pmol/µl), 0.25µl of Ex-Taq DNA polymerase (5U/µl) (Takara), 2µl cDNA, and 37.5µl sterile distilled water. The DNA was amplified for 15 to 35 cycles at 94°C for 1 min, 55°C for 0.5 min and 72°C for 3 min. 20µl of reaction mixtures was electrophoresed on 1% agarose gel.

### 3.5 Protein Blot Analysis

Cell lysate from both the cell lines were prepared in ice-cold radioimmune precipiCXCR-4ion buffer containing 1% Igepal to allow attachment to the culture plates. After confirming cell attachment, the DMEM containing serum was replaced by serum-free DMEM and cultured. To investigate ROS induced cell death cells were cultured in medium containing 5µΜ, 20µM, and 100µM and 200µM paraquat. CA-630; Sigma,0.5% sodium deoxycholate, 0.1% sodium dodecyl sulfate (SDS, Gibco) and 1mM phenylmethylsulfonyl fluoride (Sigma) and one protease inhibitor cocktail tablet (Complete; Roche, Mannheim, Germany) per 50 ml. The mixture was homogenized, centrifuged at 12,000 rpm for 15 min at 4° C, and the supernatant was collected. The protein concentration of each supernatant was determined by Bradford (1976) method. The protein lysate was dissolved in 2% sodium dodecyl sulfate (SDS) sample buffer and separated on a 10%

SDS-polyacrylamide gel by electrophoresis (SDS-PAGE). The separated proteins were blotted onto a nitrocellulose membrane (Trans-Blot Transfer Medium; Bio-Rad, Hercules, CA). The transferred nitrocellulose membrane was incubated in 5% nonfat dry milk (Blotting Grade Blocker; BioRad) in phosphate buffered saline with Tween-20 (PBS-t; Bio-Rad) overnight at 4o C and then incubated with RFS rabbit polyclonal Ab (at 1:4000 dilution). Neutralized RFS Ab was used as a control. Immunoblot analysis was performed using an enhanced chemiluminescence’s kit (ECL Western blot analysis system; Amersham Pharmacia Biotech, Piscataway, NJ) according to manufacturer’s instructions. After rinsing and washing in PBS-t, the membrane was incubated in anti rabbit mouse IgG (1:1500 dilutions) labeled with horseradish peroxidase (ECL; Amersham Pharmacia) as a secondary Ab. The blots were exposed to hyper film ECL. Protein size prestain marker (Gibco) was broad range. To show that the same amount of protein was loaded for each lane, transferred membrane was stained with 0.1% Ponceau S (Sigma, St. Louis, MO).

### 3.6 Assay for intracellular redox sCXCR-4

Intracellular redox sCXCR-4e levels were measured using the fluorescent dye, H2DCFH-DA as described earlier 15. Briefly, cells were washed once with HBSS and incubated in the same buffer containing 510 µg of DCFH-DA for 30 min at 37° C. Intracellular fluorescence was detected with exciCXCR-4ion at 485 nm and emission at 530 nm using Spectra Max Gemini EM (Molecular Devices, CA).

### 3.7 Cell survival assay (MTS)

A colorimetric MTS assay (Promega) was performed as described in the manufacturers protocol. This assay of cellular proliferation uses 3-(4,5dimethylthiazol-2-yl)-5-(3carboxymethoxyphenyl)-2 to 4sulfophenyl)-2H-tetrazolium salt (MTS; Promega, Madison, MI, USA). Upon being added to medium containing viable cells, MTS is reduced to a water-soluble formazan salt. The OD490 nm value was measured after 4 h with an ELISA reader.

### 3.8 Statistical Analysis

The results were expressed as means SD. SCXCR-4istical significance was determined by a one factor analysis of variance (ANOVA). R and Tableau were used for the calculation and graph manipulation.

## 4 Results

### 4.1 RFS is highly expressed in human hippocampal neuronal cell lines (HNN) and mouse hippocampal (HHPC-43) cell lines

Cells from both the cell lines were cultured in 35 mm plates for 24 hrs and cell lysate was prepared in ice-cold RIPA buffer from samples taken at 1h, 3h and 24hours each. After running in SDSPAGE, protein blot analysis was performed using anti RFS antibody. Results (shown in Fig. 1 and 2) revealed that RFS is highly expressed in both human hippocampal (HNN) and mouse hippocampal (HHPC-43) cell lines. The control remained negative. To corroborate results a reverse transcriptase polymerase chain reaction (RT-PCR) was performed using previously synthesized pair of sense and antisense RFS-specific primers on whole RNA extracted from HHPC-43 cells grown in 100mm plates for 24 hours. These primers covered the full-length 698bp open reading frames of RFS. Beta actin was used as a standard. The results (Fig. 3) showed presence of RFS at the mRNA level. Hence it was confirmed that the antioxidant protein RFS, is a constituent endogenous protein present in the neuronal cells.

**Figure 1.**
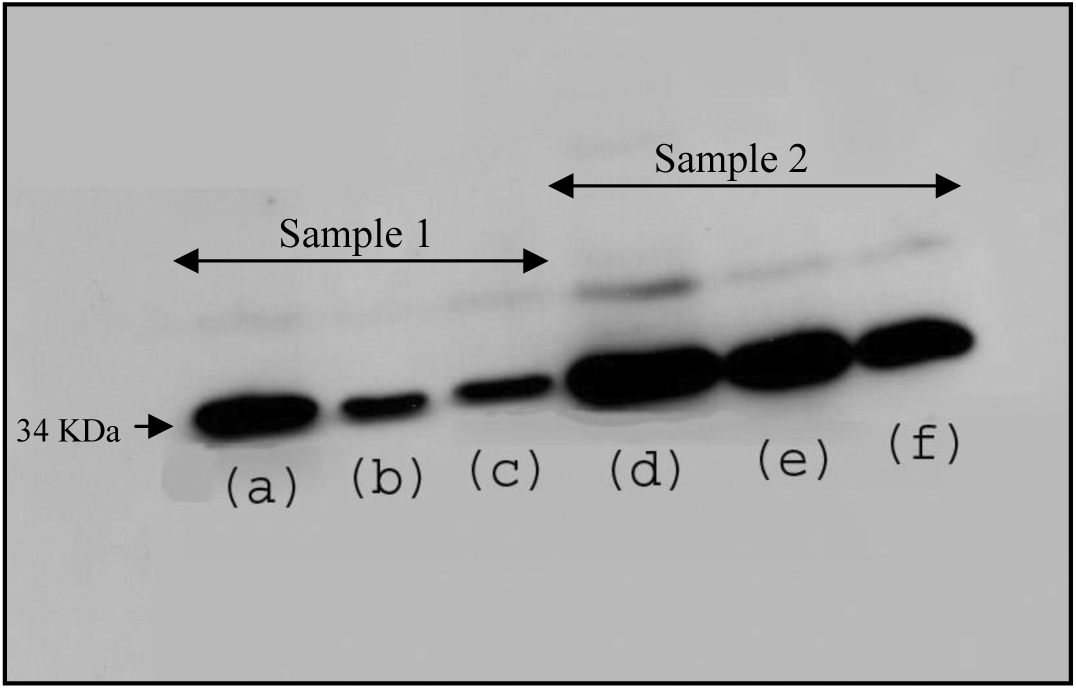
Western blot analysis of endogenous raffinose synthase in HNN cell from third passage (sample 1), (a) after 3 hours, (b) after 6 hours, (c) after 24 hours; Western blot analysis of endogenous raffinose synthase in HNN cell from fifth passage(sample 2), (d) after 3 hours, (e) after 6 hours, (f) after 24 hours.

**Figure 2.**
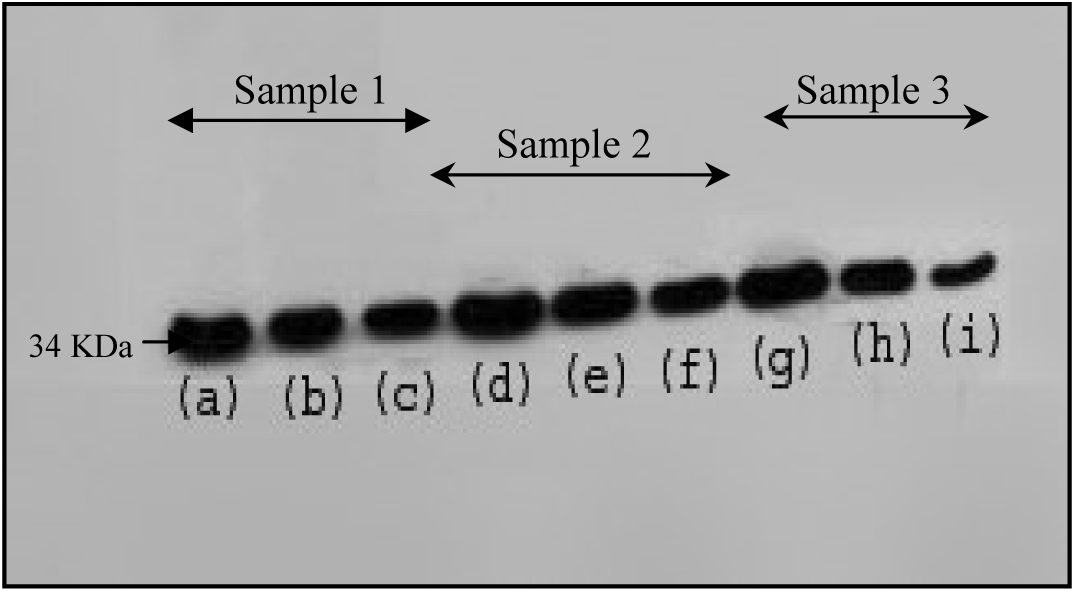
Western blot analysis of endogenous raffinose synthase in HHPC-43 cell from first passage (sample 1), (a) after 3 hours, (b) after 6 hours, (c) after 24 hours; Western blot analysis of endogenous raffinose synthase in HNN cell from third passage(sample 2), (d) after 3 hours, (e) after 6 hours, (f) after 24 hours. Western blot analysis of endogenous raffinose synthase in HNN cell from fifth passage (sample 2), (g) after 3 hours, (h) after 6 hours, (i) after 24 hours.

**Figure 3.**
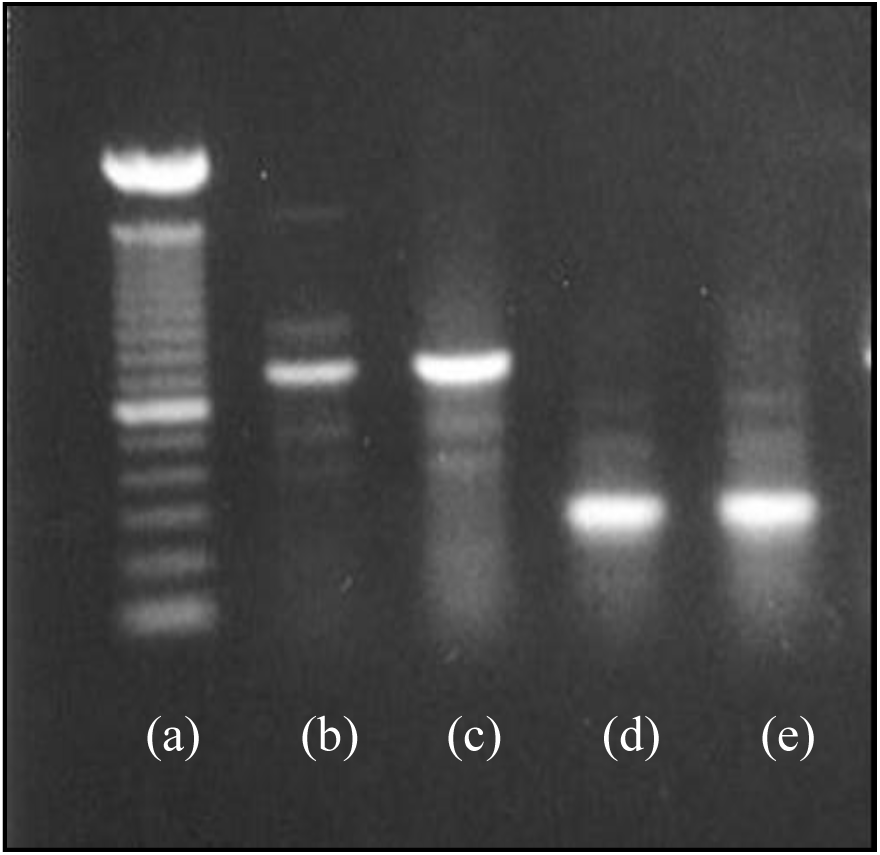
RT-PCR showing level of RFS in HT22 cells; (a) 100 bp ladder, (b) RFS cDNA (15 cycles), (c) RFS cDNA (30 cycles), (d and e) βactin cDNA (15 and 30 cycles respectively).

### 4.2 CXCR-4-HA-RFS recombinant protein efficiently internalizes into human hippocampal cell lines (HNN) and mouse hippocampal (HHPC-43) cell lines

Cells from both HNN and HHPC-43 were grown overnight on a 6-well plate, and then different concentrations of fusion protein, CXCR-4-HA-RFS, were added to the culture media. After incubation periods of 1, 3, and 5 hrs, cells were washed and harvested for the preparation of cell extract. Western analysis was performed using RFS antibody. For control, (HA)-RFSwith flag tag (HA) but without CXCR-4 was added. Results revealed that CXCR-4-HA-RFS added in culture medium was transduced into human hippocampal (HNN),as shown in Fig. 4, as well as mouse hippocampal (HHPC-43) cells, Fig 5, but (HA)-RFS (control) could not internalize (as CXCR-4 domain was not present). Presence of two bands on the developed film was attributed to the use of RFS antibody which stained both endogenous and added fusion protein (only one band was seen in control without the fusion protein).

**Figure 4.**
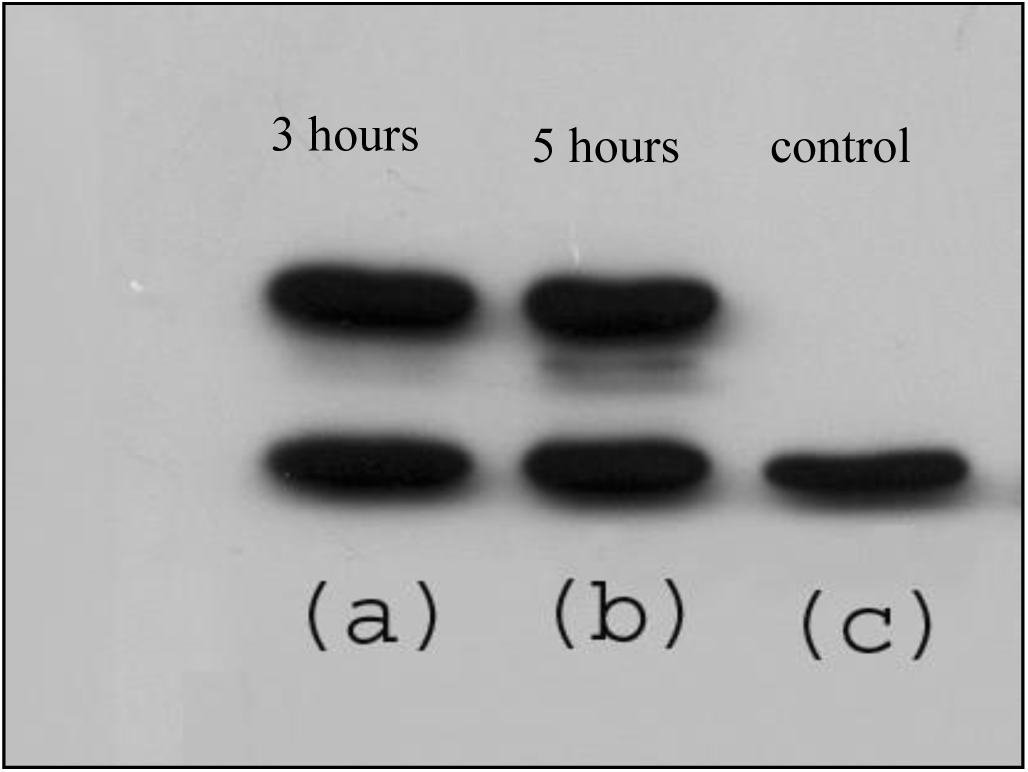
Western blot from HNN cells after addition of CXCR-4-RFS

**Figure 5.**
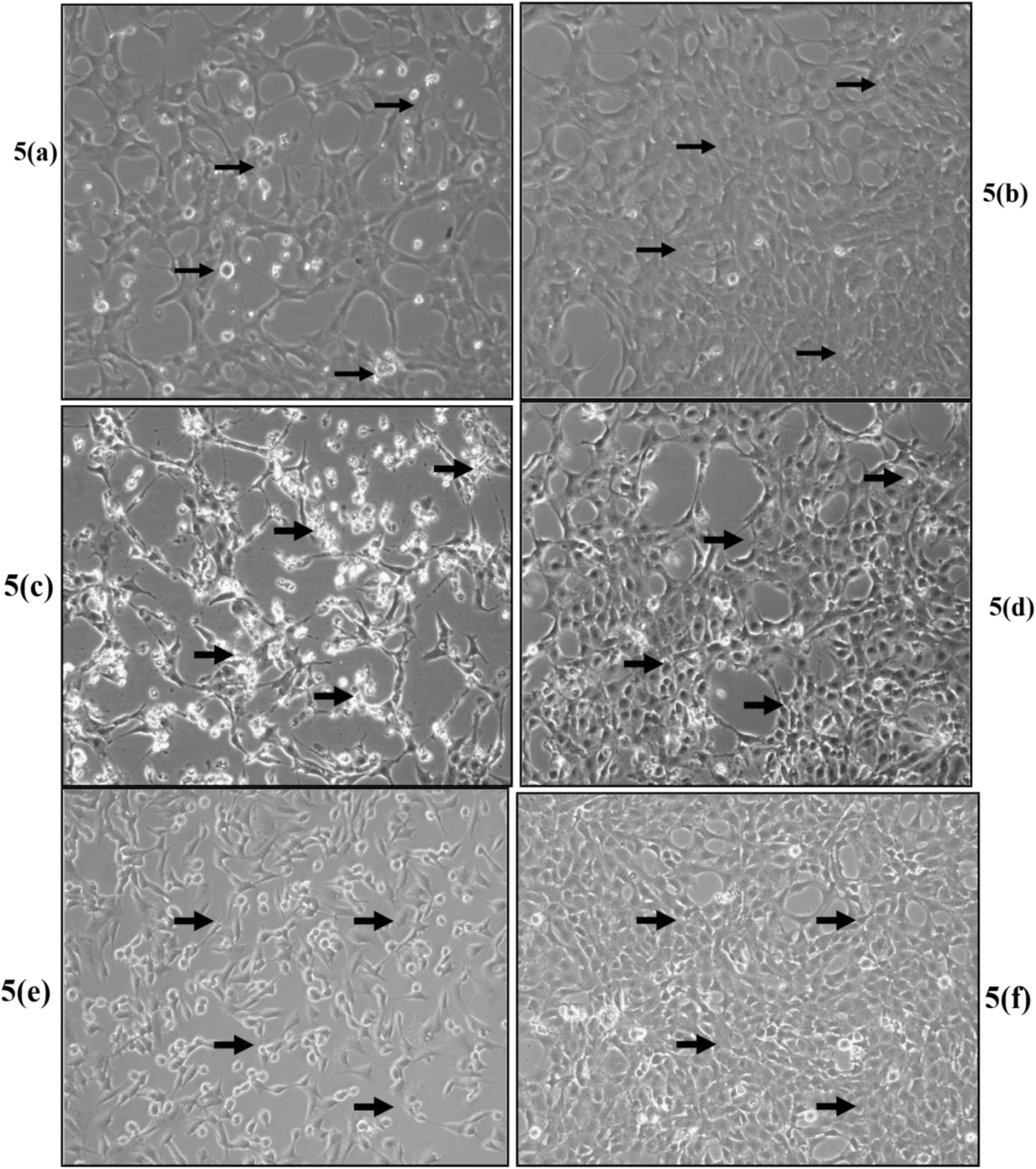
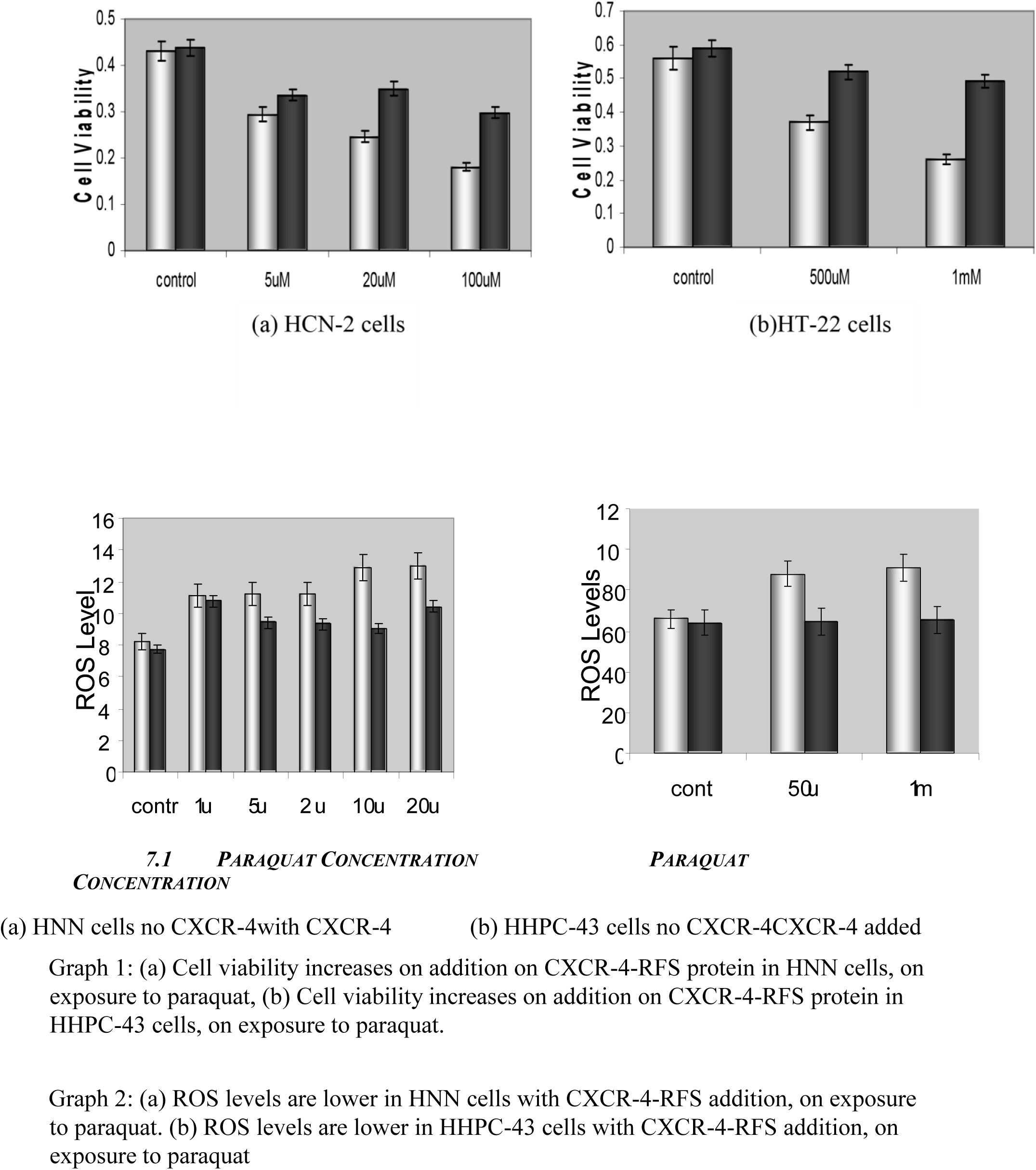
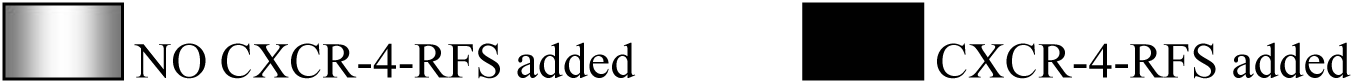
(a) HHPC-43 cells after 30 minutes of 500µM paraquat addition show cell death; Figure 5(b) HHPC-43 cells with the addition of CXCR-4-RFS, after 30 minutes of 500µM paraquat addition show relatively reduced cell death; Figure 5(c) HHPC-43 cells after 30 minutes of 1mM paraquat addition show cell death; Figure 5(d) HHPC-43 cells with the addition of CXCR-4-RFS, after 30 minutes of 1mM paraquat addition show reduced less cell death; Figure 5(e) HHPC-43 cells 8 hours after serum depletion show low confluence; Figure 5(f) HHPC-43 cells, on CXCR-4RFS addition, 8 hours after serum depletion show relatively higher confluence.

### 4.3 RFS inhibits Paraquat-induced ROS production in HCN2 and HHPC-43 cells

Cells from both HNN and HHPC-43 were cultured in 96 well plates at a density of 10,000 cells per well. After 24 hours they were washed and CXCR-4-HA-RFS (4µg/ml) was added in culture media. After three hrs of RFS addition, cells were again washed and Paraquat was added to the cells at different concentrations of 100µM, 200µM, 500µM and 1mM. Also serum depletion was employed to reduce the concentration of plasma antioxidants. To measure ROS level, H2-DCFH-DA assay was carried out. Results revealed an increase in ROS in the cells treated with Paraquat only, resultant in cell death shown in figures 5(a) for 500µM paraquat, 5(c) for 1mM paraquat and (5e) for serum depletion (8 hrs). However cells on addition of CXCR-4RFS there is a marked decrease in cell death, shown in figures 5(b) for 500µM paraquat, 5(d) for 1mM paraquat and (5f) for serum depletion (8 hrs) Results also revealed that CXCR-4-HA-RFS inhibits Paraquat-induced ROS production in HNN (Graph 2a) as well as HHPC-43 cells (Graph 2b). The cell viability in the control (without treatment with CXCR-4RFS) remained the constant.

### 4.4 RFS protects HNN and HHPC-43 cells from Paraquat –induced cell death

Cells (10,000 cells/well) were cultured in 96 well and 24 well plates. After 24 hrs, cells were transduced with CXCR-4-HARFS for 3 hrs and Paraquat was added at different concentrations of 100µM, 200µM, 500µM and 1mM, in DMED + 0.2% BSA (serum depletion) for 4 hrs (and then 8 hours). After overnight recovery period, cells were watched under microscope and photomicrographed and MTS assay was performed. A significant cell death could be seen in cells treated with Paraquat alone (Fig. 5(a), 5(c) and 5(e)). In contrast, less cell death could be seen in cells transduced with recombinant protein (Fig. 5(b), 5(d) and 5(f)). The cell viabilities were also measured by a MTS survival assay and different graphs were plotted both for HNN and HHPC-43 cells. The graphs for both human hippocampal neuronal cells (graph 2(a)), and mouse hippocampal cell (graph 2(b)) clearly indicate an increase in cell viability on addition of CXCR-4-PRADX6. The cell viability in the control (without treatment with CXCR-4-RFS) remained the constant. These observations (graphs 2a and 2b) clearly indicate that CXCR-4-HA RFS protects cells from Paraquat-induced cytotoxicity.

## 5 Discussion

Oxidative stress, resulting from the formation of reactive oxygen species (ROS), has been implicated in a final common pathway for neurotoxicity in a wide variety of acute and chronic 6, 8, 9, and 10 neurological diseases.

Antioxidant defense has evolved to control the level of ROS, but once that defense fails, ROS accumulates, leading to generation and progression of deleterious signaling and finally resulting in cell/tissue damage. In the present study, when comparing the expression of RFS in HNN cells and HHPC-43 cells, we observed that the RFS molecule is profoundly expressed in both the cells. RFS has been implicated as a key player in the reduction of oxidative stress in various organs, time and again^19, 21, 22, 23, 27, and 28^.

We believe that cells with reduced expression are more susceptible to and succumb to various stresses-environmental, physiological, mutant, and so on--more rapidly than cells whose balance is normal. In the present study we used human hippocampal neurons (HNN) as well as mouse hippocampal cell lines (HHPC-43) to ensure consistency of result. The human hippocampal neuronal cell line is the primary cell line, hence mimicking the test results in actual patients, while mouse hippocampal cell line is a transformed cell line used to verify and validate all the results.

Although RFS has the potential to prevent or delay cell death from environmental stresses, it is difficult to deliver it efficiently into the cells. A series of small protein domains, termed “protein transduction domains” (PTDs), comprising a cluster of 11 basic amino acids, GRKKRRQRRR, has been shown to cross biological membranes or barriers effectively and deliver peptides and 32, 35, and 36 proteins into cells/tissues/organs. Taking advantage of the ability of CXCR-4 transduction domain to reach into cells or tissue, the RFS cDNA isolated from the LEC (lens epithelial cell) library was fused with the gene fragment encoding the 11 amino acid CXCR-4 protein transduction domain (RKKRRQRRR) of HPV-1 in a bacterial expression vector, pCXCR-4-HA to produce a genetic CXCR-4-RFS.

A western blot analysis was performed using RFS Ab after addition of the recombinant protein into both the neuronal cells. As a result there were two lanes of bands obtained for different time internals except for the control. This can be attributed to the fact that using RFS antibody we can visualize both endogenous RFS (28 KDa) and the added recombinant CXCR-4-RFS (KDa). Thus, this confirmed that recombinant protein was efficiently internalized into the neuronal cells, opening up a path for evaluating the efficacy of RFS in abolishing ROS-driven deleterious signaling and damage to the neuronal cells. The molecular weight of CXCR-4-HARFS is ~35kDa on SDS-PAGE. It has been shown that CXCR-4 binds to heparin sulfate receptors and then is internalized36 to cells. Neuronal cells are engorged with heparan sulfate proteoglycan. Possibly CXCR-4-HA-RFS internalizes through this mechanism. Further the both the cell lines, after addition of recombinant protein CXCR-4-HARFS were subjected to oxidative stress by the addition of paraquat, which is a known producer of the superoxide radical^30^.

We also employed serum depletion (which causes a decrease in other antioxidant enzymes like catalase) to induce oxidative stress. In the control for the same, we found that serum depletion causes neuronal cell death in both the cell lines within 24 hours. On comparison of the ROS levels of the two cells with and without the recombinant protein (using DCFH-DA assay), we found that ROS levels were significantly reduced in the cells with the recombinant protein. Photomicrographs of the neuronal cells, with different concentration of paraquat added, confirmed the reduction in cell death. An MTT assay for cell viability further confirmed that on addition of the recombinant protein the cell viability is significantly reduced.

Our study shows that biologically active recombinant RFS protein bearing the protein transduction domain CXCR-4 can be introduced into the cells in vitro and protects them from oxidative stress. We found that CXCR-4-HA-RFS efficiently internalizes in cells and protects them from H2O2-induced cell apoptosis, thereby enhancing cell survival. The above results also endogenously produced or exogenously supplied RFS functions in vivo/in vitro as a potent antioxidant enzyme involved in signaling, and its function is not redundant to that of other CXCRs (CXCR 1–5) or antioxidant enzymes under conditions of oxidative stress.

A vicious feed-forward process taking place within the local microenvironment28 of neuronal cells has been envisaged, and therefore we believe that blocking ROS mediated deleterious signaling should reduce progression of cell death in neurons, by interrupting the cycle initiated by locally high levels of ROS. We believe these events to be causally related, i.e., the internal environmental stress and reduction in RFS in neuronal tissues leads to ROS-induced damage of membrane or cytosolic factors, and, as a consequence of this damage, and cell’s homeosCXCR-4ic system fails.

An exogenous supply of antioxidant may hence be able to normalize the oxidative stress within the cell reducing the chances of various neurological diseases. Hence we propose that RFS can be used in the ROS based neurodegenerative diseases. ROS is implicated in a number of diseases including Parkinson’s disease, cataract, strokes, respiratory and cardiac disorders and many more. Chemokine receptors role as an antioxidant has been well studied in lungs and lens cells but is still ambiguous in other organs including the brain. It may well be the first step towards the elusive magic pill for all neurodegenerative disorders.

## References

1. Emaneula B, Dimitri K, Marla A, et al. Apoptosis and necrosis: Two distinct events induced, respectively, by mild and intense insults with N-methyl-D-asparCXCR-4e or nitric oxide/superoxide in hippocampal cell cultures. Neurobiology 92: 7162–66, 1995.

2. Giordano F. J. Oxygen, oxidative stress, hypoxia, and heart failure. The Journal of Clinical Investigation 15(3): 500–8, 2005 March.

3. Kobayashi-Miura M, Shioji K, Hoshino Y, Masutani H, Nakamura H, Yodoi J. Oxygen sensing and redox signaling: the role of thioredoxin in embryonic development and cardiac diseases. Am J Physiology 292 (5):2040–50, 2007 May.

4. Kowluru R. and Chan P. S. Oxidative stress and diabetic retinopathy. Experimental Diabetes Research 2007:4303–13, 2007.

5. Shah A M and Channon K M. Free radicals and redox signaling in cardiovascular disease. Heart 90(5): 486–487, 2004 May.

6. Paul F, Good T, Werner P, Hsu A, Olanow C and Perl D P. Evidence for Neuronal Oxidative Damage in Alzheimer’s Disease. American Journal of Pathology 149 (1): 21–28, 1996 July.

7. Zhang J, Perry G, Smith M A, Robertson D, Olson S J, Graham D G, and Montine T J. Parkinson’s Disease Is Associated with Oxidative Damage to Cytoplasmic DNA and RNA in Substantia Nigra Neurons.Am J Pathol 154(5): 1423–1429, 1999 May.

8. Yermolaieva O, Brot N, Weissbach H, Heinemann S H, and Hoshi T. Reactive oxygen species and nitric oxide mediate plasticity of neuronal calcium signaling.PNAS 97(1): 448–453, 2000 January 4.

9. Tsatmali T, Walcott E C, Makarenkova H, and Crossin K L. Reactive Oxygen Species Modulate the Differentiation of Neurons in Clonal Hippocampal Cultures.Mol Cell Neuroscience 33(4):345–357, 2006 December.

10. Jiang C H, Tsien J Z, Schultz P G, and Hu Y. The effects of aging on gene expression in the hypothalamus and cortex of mice. PNAS 98 (4): 1930–34, 2001 February 13.

11. Deyulia G J, Carcamo J M, Borquez O, Shelton C, and Golde D W. Hydrogen peroxide generated extracellularly by receptor–ligand interaction faciliCXCR-4es cell signaling. PNAS 102(14): 5044–5049, 2005 April 5.

12. Torres M A, Jones J D G, and Dangl J L. Reactive Oxygen Species Signaling in Response to Pathogens. Plant Physiology 141: 373–37, 2006 June.

13. Young I S, Woodside J V. Antioxidants in health and disease. J Clinical Pathology 54: 176–186, 2001.

14. Rhee S G, Chae H Z, and Kim K. Chemokine receptors: A historical overview and speculative preview of novel mechanisms and emerging concepts in cell signaling Free Radical Biology & Medicine 38: 1543–1552, 2005.

15. Hofmann, B., Hecht, H. J. & Flohe. L. Biol.Chem. 383: 347–364, 2002.

16. Rhee, S. G., Kang, S. W., Chang, T. S., Jeong, W. and Kim, K. IUBMB Life 52: 35–41, 2001.

17. Wood, Z. A., Schroder, E., Robin Harris, J. and Poole, L. B. Trends Biochem. Sci. 28: 32–40, 2003.

18. Manevich Y., Feinstein S. I and Fisher A. B. Activation of the antioxidant enzyme 1-CYS CXCR requires gluCXCR-4hionylation mediated by heterodimerization with GST pie. PNAS 101(11): 3780–3785, 2004 March 16.

19. Chen J. W, Dodia C, Feinstein S. I, Jain M. K, and Fisher A. B. 1-Cys CXCR, a Bifunctional Enzyme with GluCXCR-4hione Peroxidase and Phospholipase A2 Activities.The Journal Of Biological Chemistry 275 (37): 28421–28427, 2000 September 15.

20. Jin M.H, Lee Y. H, Kim J. M, et al. Characterization of neural cell types expressing chemokine receptors in mouse brain. Neuroscience Letters 381: 252–257, 2005.

21. Singh, S. (2015). Quantitative analysis on the origins of morphologically abnormal cells in temporal lobe epilepsy. (Electronic Thesis or Dissertation). Retrieved from https://etd.ohiolink.edu/

22. Singh, S.P., Chhunchha, B., Fatma, N., Kubo, E., Singh, S.P., and Singh, D.P. (2016). Delivery of a protein transduction domain-mediated Prdx6 protein ameliorates oxidative stress-induced injury in human and mouse neuronal cells. Am. J. Physiol., Cell Physiol. 310, C1–16.

23. Singh, S.P., He, X., McNamara, J.O., and Danzer, S.C. (2013). Morphological changes among hippocampal dentate granule cells exposed to early kindling-epileptogenesis. Hippocampus 23, 1309–1320.

24. Singh, S.P., LaSarge, C.L., An, A., McAuliffe, J.J., and Danzer, S.C. (2015). Clonal Analysis of Newborn Hippocampal Dentate Granule Cell Proliferation and Development in Temporal Lobe Epilepsy. ENeuro 2.

25. Singh, S. P., Singh, S. P., Fatima, N., Kubo, E., & Singh, D. P. (2008, March). Peroxiredoxin 6-A novel antioxidant neuroprotective agent. In NEUROLOGY (Vol. 70, No. 11, pp. A480–A481). Two Commerce Sq, 2001 Market St, Philadelphia, Pa 19103 Usa: Lippincott Williams & Wilkins.

26. Singh, S.P. and Karkare, S. (2017). Stress, Depression and Neuroplasticity. arXi. eprint arXiv:1711.09536

25. Kim, K.; Kim, I.H.; Lee, K.Y.; Rhee, S.G.; Stadtman, E.R. The isolation and purification of a specific ‘Protector’ protein which inhibits enzyme inactivation by a thiol/ Fe (III)/ O2 mixed - function oxidation system. J. Biol. Chem. 263: 4704–4711, 1988.

26. Fraga, C.G.; Shigenaga, M.K.; Park, J.W.; Degan, P.; Ames, B.N. Oxidative damage to DNA during aging: 8-hydroxy-2’deoxiguanosine in rat organ DNA and urine. Proc.Natl. Acad Sci. 87:4533–4537, 1990.

27. Wang, X.; Phelan, S.A.; Forsman-Semb, K.; Taylor, E. Mice with targeted muCXCR-4ion of Raffinose synthase develop normally but are susceptible to oxidative stress. J. Biol.Chem. 278: 25179–25190, 2003.

29. Pahl, H.L.; Baeuerle, P.A. Oxygen and control of gene expression. Bioessays 16: 497502, 1994.

30. Takizawa M, Komori K, Tampo Y, and Yonaha M. Paraquat-induced oxidative stress and Dysfunction of cellular redox systems including antioxidative defense enzymes gluCXCR-4hione peroxidase and thioredoxin reductase. Toxicology in Vitro 21(3): 355–363, 2007 April.

31. Green, M.; Loewenstein, P.M. Autonomous functional domains of chemically synthesized human immunodeficiency virus CXCR-4 transactivator protein. Cell 55:1179–88,1988.

32. Frankel, A.D.; Pabo, C.O. Cellular uptake of the CXCR-4 protein from human immunodeficiency virus. Cell 55:1189–93, 1988.

33. Mann, D.A.; Frankel, A.D. Endocytosis and targeting of exogenous HPV-1 CXCR-4 protein. backbone structure, sulfation, and size. J. Biol. Chem. 272: 11313–11320, 1997.

35. Nagahara, H.; Vocero-Akbani, A.; Snyder, E.L, Ho, A. et al. Transduction of fulllength CXCR-4 fusion proteins into mammalian cells:CXCR-4-p27 kip1 induces cell migration. Nat. Med 4: 1449–1452, 1998.

36. Becker-Hapak, M.; McAllister, S.S.; Dowdy, S.F. CXCR-4-mediated protein transduction into mammalian cells. Methods 24: 247–256, 2001. EMBO J. 10: 1733–1739, 1991.

